# Generation of Recombinant Rotavirus Expressing NSP3-UnaG Fusion Protein by a Simplified Reverse Genetics System

**DOI:** 10.1101/678292

**Authors:** Asha A. Philip, Jacob L. Perry, Heather E. Eaton, Maya Shmulevitz, Joseph M. Hyser, John T. Patton

**Author notes:** **Corresponding address:** John T. Patton, Department of Biology, Indiana University, 212 S. Hawthorne Drive, Simon Hall 011, Bloomington, IN 47405 USA. **Email addresses**: Asha A. Philip,; Heather E. Heaton,; Maya Shmulevitz,; John T. Patton.

## Abstract

Rotavirus is a segmented double-stranded (ds)RNA virus that causes severe gastroenteritis in young children. We have established an efficient simplified rotavirus reverse genetics (RG) system that uses eleven T7 plasmids, each expressing a unique simian SA11 (+)RNA, and a CMV support plasmid for the African swine fever virus NP868R capping enzyme. With the NP868R-based system, we generated recombinant rotavirus (rSA11/NSP3-FL-UnaG) with a genetically modified 1.5-kB segment 7 dsRNA that encodes full-length NSP3 fused to UnaG, a 139-aa green fluorescent protein (FP). Analysis of rSA11/NSP3-FL-UnaG showed that the virus replicated efficiently and was genetically stable over 10 rounds of serial passage. The NSP3-UnaG fusion product was well expressed in rSA11/NSP3-FL-UnaG-infected cells, reaching levels similar to NSP3 in wildtype rSA11-infected cells. Moreover, the NSP3-UnaG protein, like functional wildtype NSP3, formed dimers *in vivo*. Notably, NSP3-UnaG protein was readily detected in infected cells via live cell imaging, with intensity levels ~3-fold greater than that of the NSP1-UnaG fusion product of rSA11/NSP1-FL-UnaG. Our results indicate that FP-expressing recombinant rotaviruses can be made through manipulation of the segment 7 dsRNA without deleting or interrupting any of the twelve open reading frames of the virus. Because NSP3 is expressed at levels higher than NSP1 in infected cells, rotaviruses expressing NSP3-based FPs may be a more sensitive tool for studying rotavirus biology than rotaviruses expressing NSP1-based FPs. This is the first report of a recombinant rotavirus containing a genetically engineered segment 7 dsRNA.

**Importance:** Previous studies have generated recombinant rotaviruses that express fluorescent proteins (FPs) by inserting reporter genes into the NSP1 open reading frame (ORF) of genome segment 5. Unfortunately, NSP1 is expressed at low levels in infected cells, making viruses expressing FP-fused NSP1 less than ideal probes of rotavirus biology. Moreover, FPs were inserted into segment 5 in such a way as to compromise NSP1, an interferon antagonist affecting viral growth and pathogenesis. We have identified an alternative approach for generating rotaviruses expressing FPs, one relying on fusing the reporter gene to the NSP3 ORF of genome segment 7. This was accomplished without interrupting any of the viral ORFs, yielding recombinant viruses likely expressing the complete set of functional viral proteins. Given that NSP3 is made at moderate levels in infected cells, rotavirus encoding NSP3-based FPs should be more sensitive probes of viral infection than rotaviruses encoding NSP1-based FPs.

## Introduction

Group A rotavirus (RVA) is a primary cause of acute gastroenteritis in infants and children under 5 years of age (Crawford et al., 2017). The RVA genome consists of eleven segments of double-stranded (ds)RNA, with a total size of 18.5 kbp, and is contained within a non-enveloped icosahedral capsid composed of three concentric protein layers (Settembre et al., 2011). During replication, the genome segments are transcribed, producing eleven 5’-capped, but non-polyadenylated (+)RNAs (Imai et al., 1983; Trask et al., 2012). The (+)RNAs are generally monocistronic, each with a single open reading frame (ORF) that specifies one of the six structural (VP1-VP4, VP6-VP7) or six nonstructural (NSP1-NSP6) proteins of the virus. The (+)RNAs also serve as templates for the synthesis of dsRNA genome segments (Gugliemli et al., 2010).

Insights into RVA biology have been severely hampered by the lack of a reverse genetics (RG) system to unravel details into the replication and pathogenesis mechanisms of the virus. This limitation was recently overcome by Kanai et al. (2017), who described the development of a fully plasmid-based RG system that allowed genetic engineering of the prototypic RVA simian SA11 strain. Key to the RG system was co-transfection of baby hamster kidney cells expressing T7 RNA polymerase (BHK-T7 cells) with T7 plasmids directing synthesis of the eleven SA11 (+)RNAs, two CMV plasmids encoding the vaccinia virus D1L-D12R capping enzyme complex (Kyrieleis et al., 2014), and another CMV plasmid encoding the avian reovirus p10FAST fusion protein (Salsman et al., 2005). Subsequent publications described changes to the Kanai RG system designed to reduce its complexity and/or enhance the recovery of recombinant virus. Notably, Komoto et al (2018) showed that recombinant virus could be produced simply by transfecting BHK-T7 cells with eleven SA11 T7 plasmids, with the caveat that plasmids for the viroplasm building blocks (NSP2 and NSP5) (Fabbretti et al., 1999; Eichwald et al., 2004) be added at levels three-fold greater than the other plasmids. Of possible significance, the Komoto RG system used a set of SA11 T7 plasmids with vector backbones that differed in size and sequence from the SA11 T7 plasmids described by Kanai et al (2017).

Plasmid-based RG systems have been used to modify several SA11 genome segments, with the focus mostly on segment 5, which encodes NSP1 (Kanai et al., 2017, 2018; Komoto et al., 2018). Through insertion of reporter genes into the segment 5 dsRNA, recombinant RVAs have been produced that express fluorescent reporter proteins (FPs) (e.g., mCherry, eGFP) (Kanai et al., 2017, 2018; Komoto et al., 2018); these FP-RVAs are important tools for studying virus replication and pathogenesis via live cell imaging and other fluorescence-based approaches. Unfortunately, the segment 5 product NSP1 is expressed at low levels in infected cells and is subject to proteasomal degradation making FP-fused NSP1 proteins less than ideal probes of RVA biology (Martinez-Alvarez et al., 2013). Moreover, FP genes were inserted into the segment 5 dsRNA in such a way as to alter the NSP1 open reading frame (ORF), likely compromising the protein’s function as an IFN-antagonist (Barro and Patton, 2005; Davis and Patton, 2017) and, thus, impacting the virus’s biological properties.

In this study, we explored an alternative approach for making FP-RVAs, one relying on modification of the genome segment that expresses NSP3 (segment 7), a viral translation enhancer expressed at moderate levels in the infected cell that may not be required for virus replication (Montero et al., 2006; Gratia et al., 2015). In generating recombinant RVAs, we employed a simplified RG system requiring only a single support plasmid: a CMV expression vector for the African swine fever virus (ASFV) NP868R capping enzyme (Dixon et al., 2013).

Using the NP868R-based RG system, we produced a novel SA11 strain with modified segment 7 dsRNA that expressed NSP3 fused to the green fluorescent protein (FP) UnaG. This is the first RVA strain engineered to produce a FP that did not involve deleting or interrupting any of one of the 12 viral ORFs, thus yielding a recombinant virus likely expressing a complete set of functional viral proteins.

## Materials and methods

### Cell culture

Embryonic monkey kidney cells (MA104) were grown in M199 complete medium [Medium 199 (Lonza) and 1% penicillin-streptomycin (P/S) (Corning)] containing 5% fetal bovine serum (FBS) (Gibco) (Arnold et al., 2009). BHK-T7 cells were a kind gift from Drs. Ulla Buchholz and Peter Collins, Laboratory of Infectious Diseases, NIAID, NIH. BHK-T7 cells were grown in Glasgow complete medium [Glasgow minimum essential medium (G-MEM) (Lonza), 10% tryptone-peptide broth (Gibco), 1% P/S, 2% non-essential amino acids (NEAA) (Gibco), and 1% glutamine (Gibco)] containing 5% heat-inactivated FBS. Medium used to cultivate BHK-T7 cells was supplemented with 2% G418 geneticin (Invitrogen) every other passage.

### Plasmid construction

RVA (simian SA11 strain) plasmids pT7/VP1SA11, pT7/VP2SA11, pT7/VP3SA11, pT7/VP4SA11, pT7/VP6SA11, pT7/VP7SA11, pT7/NSP1SA11, pT7/NSP2SA11, pT7/NSP3SA11, pT7/NSP4SA11, and pT7/NSP5SA11 were kindly provided by Dr. Takeshi Kobayashi (Kanai et al., 2017) through the Addgene plasmid repository {https://www.addgene.org/Takeshi_Kobayashi/}. To generate the pCMV/NP868R plasmid, a DNA representing the ASFV NP868R ORF (Genbank NP_042794), bound by upstream *Xba*I and downstream *Bam*HI sites, was synthesized by Genscript and inserted into the *Eco*RV site of the pUC57 plasmid. A DNA fragment containing the NP868R ORF was recovered from the plasmid by digestion with *Not*I and *Pvu*I and ligated into the plasmid pCMV-Script (Agilent Technologies), cut with the same two restriction enzymes. pT7/NSP3-FL-UnaG was constructed by fusing a DNA fragment containing the ORF for 3xFL-UnaG to the 3’-end of the NSP3 ORF in pT7/NSP3SA11 using a Takara In-fusion HD cloning kit. The 3xFL-UnaG fragment was amplified from the reovirus S1 3xFL-UnaG plasmid (Eaton et al, 2017) using the primer pair 5’-tcatttggttgcgaagactacaaagaccatgacggtgattataaaga-3’ and 5’-catgtatcaaaatggtcattctgtggcccttctgtagct-3’ and the pT7/NSP3SA11 plasmid was amplified using the NSP3 primer pair 5’-ccattttgatacatgttgaacaatcaaatacagtgt-3’ and 5’-ttcgcaaccaaatgaatattgataattacatctctgtattaat-3’. pT7/NSP1-FL-UnaG was constructed by inserting the 3xFL-UnaG fragment into the NSP1 ORF of the segment 5 cDNA (position 1199) in pT7/NSP1SA11. For cloning, the pT7/NSP1SA11 plasmid was amplified using the NSP1 primer pair 5’-tgaagaagtgtttaatcacatgtcgcc-3’ and 5’-tttgatccatgtgattagtaaacaaactccaaa-3 and the 3xFL-UnaG insert was amplified from pT7/NSP3-R2A-FL-UnaG using the primer pair 5’-tcacatggatcaaacctacaaagaccatgacggtgattataaagatcat-3’ and 5’-ttaaacacttcttcatcattctgtggcccttctgtagc-3’. Transfection quality plasmids were prepared commercially (www.plasmid.com) or using a QIAprep spin miniprep kit. Primers and sequencing services were provided by Eurofin Genomics. Sequences of recombinant RVAs were determined from cDNAs prepared from viral RNAs using a Superscript III reverse transcriptase kit (Invitrogen).

### Optimized RVA RG protocol

On Day 0, freshly confluent monolayers of BHK-T7 cells were disrupted using trypsin-versene and resuspended in G418-free Glasgow complete medium containing 5% FBS. Cell numbers were determined with a Nexcelom Cellometer AutoT4 counter. The cells were seeded in the same medium into 12-well plates (2-4 x 10_5_ cells/well). On Day 1, plasmid mixtures were prepared that contained 0.8 μg each of the 11 RVA pT7 plasmids, except pT7/NSP2SA11 and pT7/NSP5SA11, which were 2.4 μg each. Included in plasmid mixtures was 0.8 μg of pCMV/NP868R. The plasmid mixtures were added to 100 μl of pre-warmed (37°C) Opti-MEM (Gibco) and mixed by gently pipetting up and down. Afterwards, 25 μl of TransIT-LTI transfection reagent (Mirus) was added, and the transfection mixtures gently vortexed and incubated at room temperature for 20 min. During the incubation period, BHK-T7 cells in 12-well plates were washed once with Glasgow complete medium, and then 1 ml of SMEM complete medium [MEM Eagle Joklik (Lonza), 10% TBP, 2% NEAA, 1% P/S, and 1% glutamine)] was placed in each well. The transfection mixture was added drop-by-drop to the medium in the wells and the plates returned to a 37°C incubator. On Day 3, 2 x10_5_ MA104 cells in 250 μl of M199 complete medium were added to wells, along with trypsin to a final concentration of 0.5 μg/ml (porcine pancreatic type IX, Sigma Aldrich). On Day 5, cells in plates were freeze-thawed 3-times and the lysates placed in 1.5 ml microfuge tubes. After centrifugation at 500 x g for 10 min (4°C), 300 μl of the supernatant was transferred onto MA104 monolayers in 6-well plates containing 2 ml of M199 complete medium and 0.5 μg/ml trypsin. The plates were incubated at 37°C for 7 days or until complete cytopathic effects (CPE) were observed. Typically, complete CPE was noted at 4-6 days p.i. for wells containing replicating RVA.

### Analysis of recombinant viruses

RVAs were propagated in MA104 cells in M199 complete medium containing 0.5 μg/ml trypsin. Viruses were isolated by plaque purification and titered by plaque assay or fluorescence focus assay on MA104 cells (Arnold et al., 2009). To determine peak viral titers, virus isolates were grown in parallel in MA104 cell monolayers until cytopathic effects reached 100%. Plaque titers in lysates were calculated from six independent plaque assays. Viral RNAs were recovered from clarified infected-cell lysates by Trizol extraction, resolved by electrophoresis on Novex 8% polyacrylamide gels (Invitrogen), and detected by staining with ethidium bromide.

For immunoblot assays, proteins in lysates prepared at 8 h p.i. from MA104 cells infected with RVA at an MOI of 5 were resolved by electrophoresis on Novex linear 8-16% polyacrylamide gels and transferred to nitrocellulose membranes. After blocking with phosphate-buffered saline containing 5% nonfat dry milk, blots were probed with guinea pig polyclonal anti-NSP3 (Lot 55068, 1:2000) or anti-VP6 (Lot 53963, 1:2000) antisera (Arnold et al., 2012), or mouse monoclonal Flag M2 (Sigma F1804, 1:2000) or rabbit monoclonal PCNA (13110S, Cell Signaling Technology (CST), 1:1000) antibody. Primary antibodies were detected using 1:10,000 dilutions of horseradish peroxidase (HRP)-conjugated secondary antibodies: horse anti-mouse IgG (CST), anti-guinea pig IgG (KPL), or goat anti-rabbit IgG (CST). Signals were developed using Clarity Western ECL Substrate (Bio-Rad) and detected using a Bio-Rad ChemiDoc imaging system. Image J analysis was used to determine the intensity of bands on immunoblots {https://imagej.net/ImageJ}.

### Assessment of genetic stability

MA104 cell monolayers in 6-well plates were infected with recombinant RVA at an MOI of ~0.1. When cytopathic effects were complete (4-5 days p.i.), the cells were freeze-thawed twice in their medium, and lysates were centrifuged at low speed to remove debris. Virus in clarified lysates was serially passaged 10-times, infecting MA104 cells with 2 μl of lysate combined with 2 ml of fresh M199 complete medium. To analyze viral dsRNA content, 0.6 ml of clarified lysates were incubated with 1 μl of RNase T1 (Fermentas, 1000 U/ml) for 30 min at 37°C, extracted with Trizol, and pelleted by ethanol precipitation. Purified dsRNA was resolved by electrophoresis on 8% polyacrylamide gels and detected by staining with ethidium bromide. Plaque purification was used to recover six virus isolates from the passage 10 pool of rSA11/NSP3-FL-UnaG. The segment 6 (VP6) and 7 (NSP3) RNAs of three isolates were sequenced by Eurofins Genomics.

### Live-cell imaging of UnaG fluorescence in infected cells

MA104 cells were seeded at a density of 25,000 cells per well in a 96-well optical bottom plate (Greiner Bio-One) and grown for 2 days at 37°C/5% CO_2_ before being infected with rSA11/NSP1-FL-UnaG or rSA11/NSP3-FL-NSP3-UnaG at ~4 fluorescent focus units per cell. After 1 h the inoculum was removed and 200 μl FluoroBrite (Thermo Fisher Scientific) was added to each well for maintenance media. The plate was mounted into an Oko Labs stage-top environmental chamber equilibrated to 37°C/5% CO_2_. Wells were imaged with a widefield epifluorescence Nikon TiE inverted microscope using a SPECTRAX LED light source (Lumencor) and a 20x Plan Fluor (NA 0.45) phase contrast objective. Fluorescence light images were recorded using an ORCA-Flash 4.0 sCMOS camera (Hamamatsu), and Nikon Elements Advanced Research v4.5 software was used for multipoint position selection, data acquisition, and image analysis. Single cells were selected as Regions of Interest (ROI) and fluorescence intensity measured for the experiment.

Fluorescence intensity values were exported to Microsoft Excel and normalized to the initial baseline fluorescence value. The baseline and maximum fluorescence values for each ROI selected were calculated and from this the change in fluorescence (ΔF) was calculated by subtracting the maximum and baseline values. The ΔF was plotted as percent change and biostatistical analyses was performed using GraphPad Prism (version 8.1) with results presented as mean ± standard deviation. Comparisons used a nonparametric Mann-Whitney test and differences were considered statistically significant for p < 0.05.

### Nucleotide sequence accession numbers

pCMV/NP868R, MH212166; pT7/NSP3-FL-UnaG, MK868472; and, pT7/NSP1-FL-UnaG, MH197081.

## Results and Discussion

### NP868R-based RG system

A previous study showed that addition of a CMV plasmid expressing the ASFV NP868R capping enzyme to the mammalian reovirus RG system significantly increased recovery of recombinant virus (Eaton et al., 2017). Based on this finding, we constructed a similar CMV plasmid for NP868R (pCMV/NP868R) and evaluated whether its co-transfection with SA11 pT7 plasmids into BHK-T7 cells was sufficient to allow recovery of recombinant RVAs. In our RG experiments, two different types of pT7 vectors for segment 7 (NSP3) were used, one (pT7/NSP3SA11) designed to introduce a wild type segment 7 dsRNA into recombinant RVA and the second (pT7/NSP3-FL-UnaG) designed to test the possibility of introducing a modified segment 7 RNA that expressed NSP3 with a fluorescent tag. To construct pT7/NSP3-FL-UnaG, the C-terminus of the NSP3 ORF in pT7/NSP3SA11 was fused to the ORF for UnaG, a 139-amino acid (aa) green fluorescent protein of the Unagi eel that utilizes bilirubin as a fluorophore (Kumagai et al., 2013) (Fig. 1). To further ease detection of the protein product of the modified pT7 plasmid, a 3x Flag sequence was inserted between the NSP3 ORF and UnaG ORF. As a result of the addition of Flag and UnaG sequences, the pT7/NSP3-FL-UnaG vector expresses a 1.6-kb RNA that encodes a 477-aa protein instead of the 1.1-kb RNA and 315-aa protein of pT7/NSP3SA11 (Fig. 1). The NSP3-FL-UnaG fusion protein retained the same RNA-binding domain, coiled-coil dimerization domain, and eIF4G-binding domain present in wildtype NSP3 (Deo et al., 2002; Groft et al., 2002).

**Figure 1.**
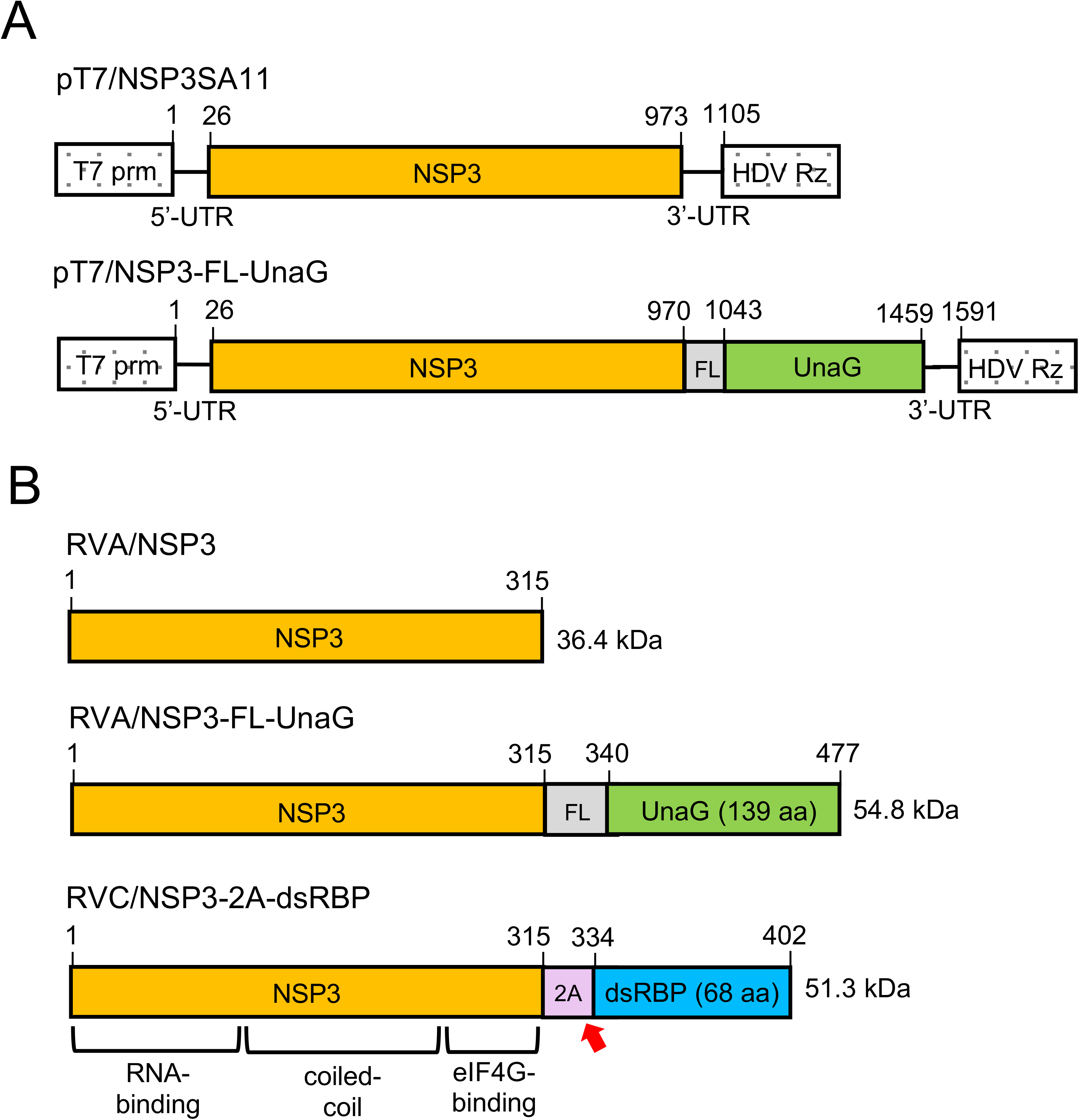
Wildtype and modified NSP3 proteins encoded by pT7 plasmids and rotaviruses. (A) Organization of pT7 plasmids expressing wild type NSP3 and NSP3-FL/UnaG (+)RNAs, indicating locations of T7 promoter (prm) and Hepatitis delta virus (HDV) self-cleaving ribozyme (Rz). Nucleotide positions are labeled. (B) Products of recombinant RVAs expressing wild type NSP3 and NSP3-FL-UnaG and RVC (Bristol strain) expressing NSP3-2A-dsRBP, including approximate locations of functional domains in NSP3 (Deo et al., 2002; Groft and Burley, 2002). Red arrow indicates the position of the stop-restart cleavage site in the 2A-like element in the RVC NSP3-2A-dsRBP ORF. Amino acid positions are labeled. FL represents a 3x Flag tag.

Rotavirus RG experiments were performed using stocks of BHK-T7 cells maintained in Glasgow medium enriched with FBS, tryptone-peptide broth, and non-essential amino acids. Mixtures of RG plasmids were transfected into BHK-T7 cells that were ~90% confluent and had been seeded into 12-well plates the day before. In our optimized protocol, plasmid mixtures included pCMV/NP868R, 3x levels of pT7/NSP2SA11 and pT7/NSP5SA11, either pT7/NSP3SA11 or pT7/NSP3-FL-UnaG, and the remaining SA11 pT7 plasmids. Transfected BHK-T7 cells were over seeded with MA104 cells to promote amplification of recombinant viruses. At 5 days post transfection, the cells were freeze-thawed thrice and large debris removed by low speed centrifugation. Recombinant viruses in cell lysates were amplified by passage in MA104 cells, plaque isolated, and amplified again in MA104 cells.

### Recombinant SA11 expressing fused NSP3-UnaG

Analysis of the cell lysates showed that transfection of BHK-T7 cells with RG plasmid mixtures using the optimized protocol supported the generation of recombinant RVAs, including SA11 isolates with a wildtype segment 7 (NSP3) dsRNA (rSA11/wt) or a modified segment 7 RNA (rSA11/NSP3-FL-UnaG) (Fig. 2). The identity of the modified segment 7 dsRNA in rSA11/NSP3-FL-UnaG was verified by gel electrophoresis (Fig. 2A), which revealed that the wildtype 1-kB segment 7 dsRNA had been replaced with a segment co-migrating with the 1.5-kB segment 5 (NSP1) dsRNA. The authenticity of the segment 7 dsRNA in rSA11/NSP3-FL-UnaG was confirmed by RT-PCR and sequencing (data not shown). Immunoblot analysis with anti-NSP3 and anti-FLAG antibodies of infected cell lysates indicated that segment 7 of rSA11/NSP3-FL-UnaG expressed a protein of the expected size (55 kD) for NSP3-FL-UnaG and did not express the wildtype 37-kD NSP3 (Fig. 2B). As a probe of the properties of NSP3-FL-UnaG, we examined whether the protein was able to form dimers in infected cells, as previously reported for wildtype NSP3 (Arnold et al, 2012). Indeed, electrophoretic analysis of rSA11/NSP3-UnaG-infected cell lysates treated with denaturing sample buffer at 25°C showed that the NSP3-FL-UnaG migrated as a dimer (Fig. 3), suggesting that the NSP3 coiled-coil dimerization domain retains its function in the fusion product. Under these same electrophoretic conditions, the VP6 inner capsid protein of both rSA11/NSP3-FL-UnaG and rSA11/wt formed trimers that were stable at 25°C (Fig. 3) (Clapp and Patton, 1991). Quantitation of bands appearing on immunoblots probed with anti-NSP3 and anti-VP6 antibodies indicated that the steady-state level of NSP3-FL-UnaG (normalized to VP6) in rSA11/NSP3-FL-UnaG-infected cells approximated that of the steady-state level of NSP3 in rSA11/wt infected cells (Fig. 3, lanes 4 versus 6). Thus, the segment 7 (+)RNAs of rSA11/NSP3-FL-UnaG and rSA11/wt appear to be translated with near equal efficiency.

**Figure 2.**
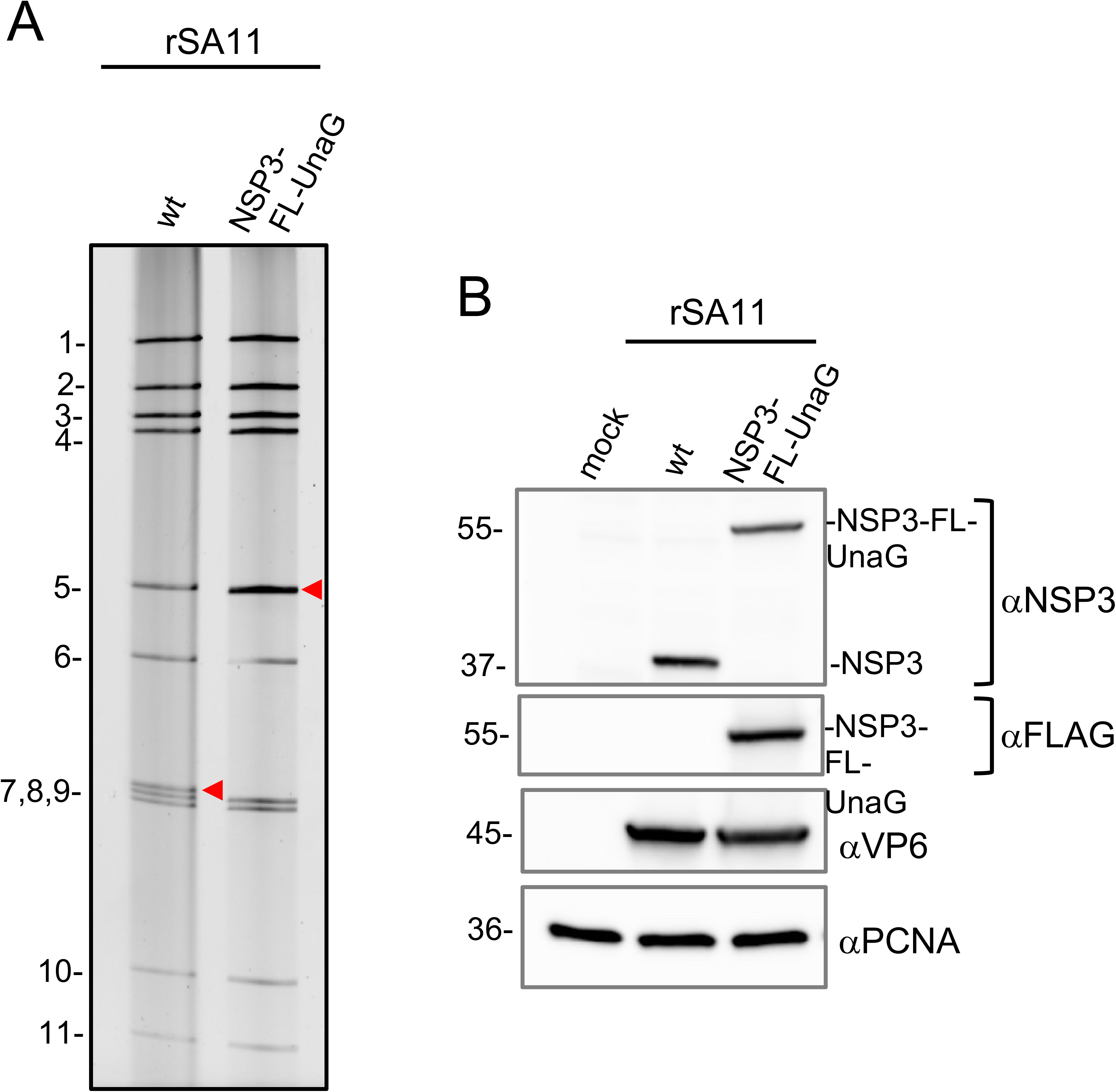
Recovery of recombinant RVA with a modified segment 7 dsRNA that expresses the fused NSP3-FL-UnaG protein. (A) Profiles of the eleven genomic dsRNAs recovered from rSA11 viruses resolved by PAGE. Red arrow notes the position of the segment 7 dsRNA. (B) Western blot analysis of proteins present at 8 h p.i. in MA104 cells infected with recombinant viruses.

**Figure 3.**
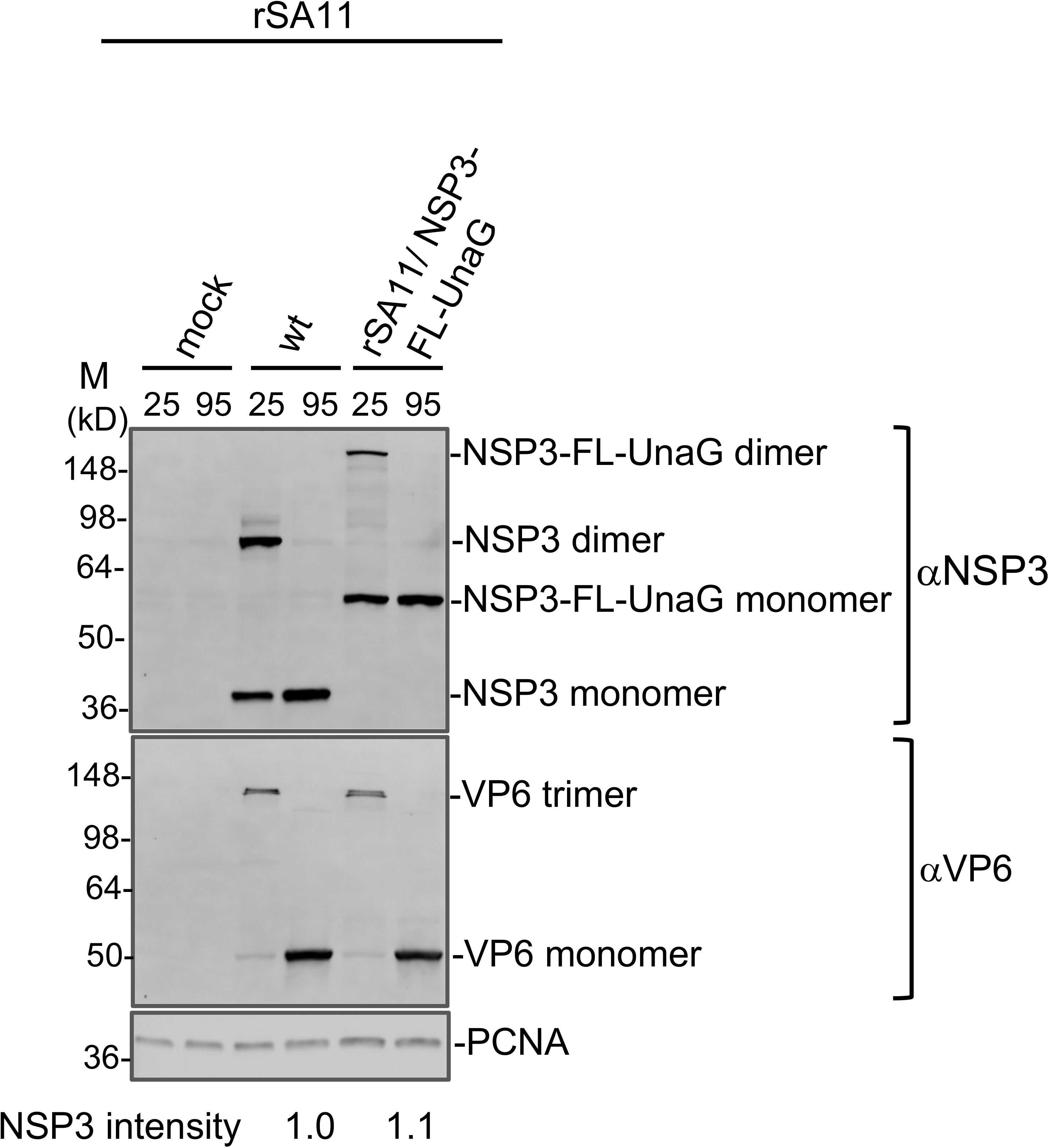
Dimerization of NSP3-FL-UnaG. MA104 cells were mock infected, or infected with rSA11/wt or rSA11/NSP3-FL-UnaG and incubated until 8 h p.i., when cells were harvested. Cell lysates were mixed with sample buffer containing sodium dodecyl sulfate and b-mercaptoethanol, incubated for 10 min either at 25 or 95°C, resolved by electrophoresis on a Novex 8-16% polyacrylamide gel, and the blotted onto a nitrocellulose membrane. Blots were probed with guinea pig polyclonal anti-NSP3 or anti-VP6 antibodies, or with a mouse anti-PCNA monoclonal antibody. Primary antibodies were detected using horseradish peroxidase-conjugated secondary antibodies. Sizes (kD) of protein markers (M) are indicated. NSP3 band intensities were determined by Image J analysis and were normalized to VP6 band intensities.

Plaque analysis showed that rSA11/wt grew to a peak titer in MA104 cells that was ~2-3-fold greater than rSA11/NSP3-FL-UnaG (1.6 - 4.8 × 10_7_ and 0.6 - 2.0 × 10_7_, respectively) and generated plaques that were ~2-fold larger (Fig. 4). Ten rounds of serial passage of rSA11/NSP3-FL-UnaG at low MOI revealed no difference in the dsRNA profile of the starting virus and the passage 10 virus, suggesting that the recombinant RVA was genetically stable (Fig. 5). To further evaluate this issue, the segment 6 (VP6) and 7 (NSP3) RNAs of three plaque isolates recovered from the passage 10 pool of rSA11/NSP3-FL-UnaG were sequenced. The sequences of segment 6 and 7 RNAs were the same as those present in the pT7/VP6SA11 and pT7/NSP3-FL-UnaG plasmids used to make the recombinant viruses. Thus, based on serial passage and sequencing of P10 viruses, rSA11/NSP3-FL-UnaG appears to stably retain its foreign sequence.

**Figure 4.**
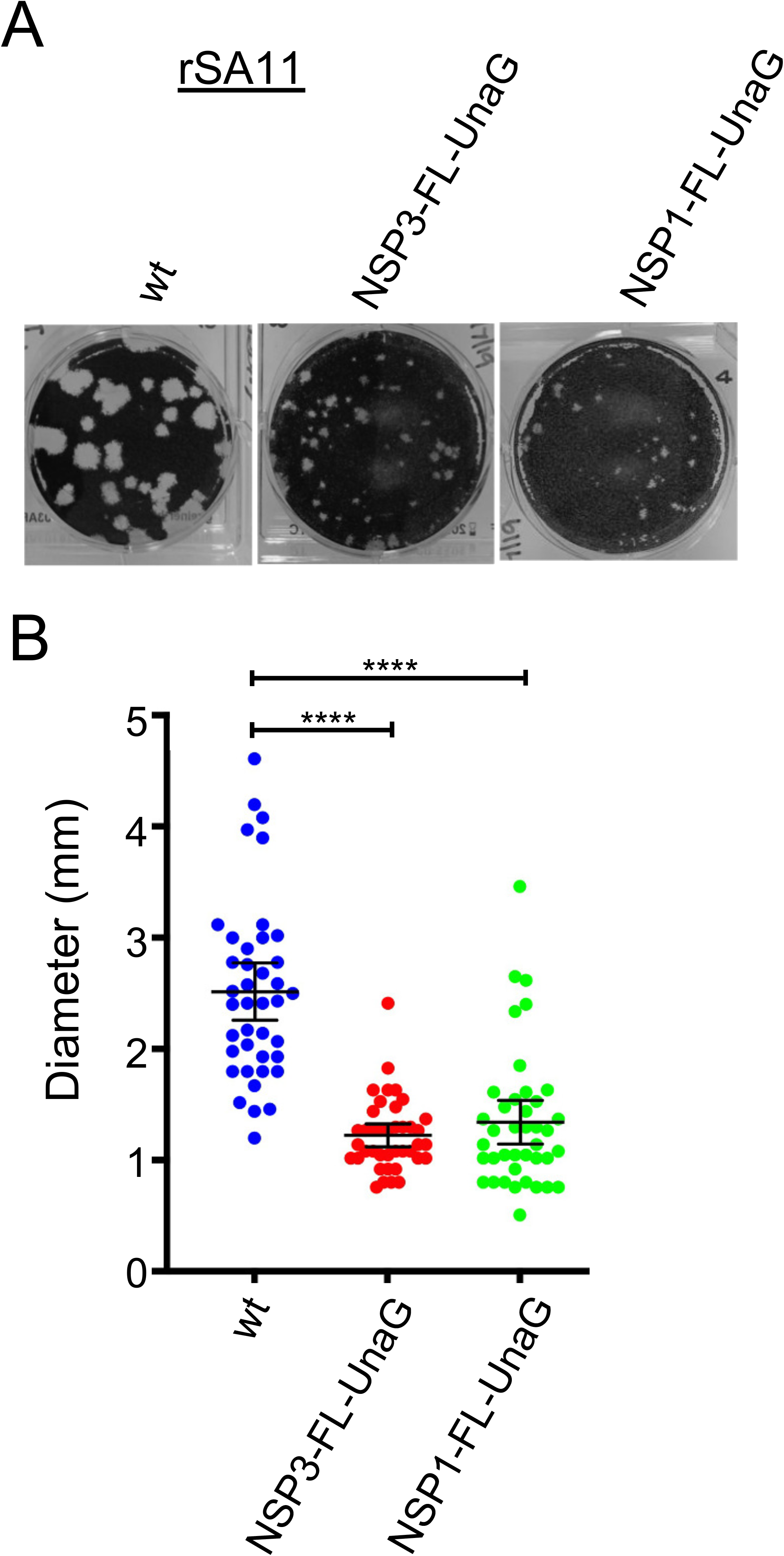
Comparison of plaques formed by wildtype rSA11 and mutant rSA11 strains containing NSP3-FL-UnaG or NSP1-FL-UnaG sequences. (A) Plaques were generated on MA104 monolayers and detected at 7 days p.i. by crystal violet staining (Arnold et al., 2009). (B) Sizes of 40 randomly selected plaques from six independent plaque assays were measured and the means determined. Mean values and 95% confidence intervals are plotted (black lines). Significance values were calculated using an unpaired Student’s t-test (GraphPad Prism, version 8). ****, *P* < 0.0001

**Figure 5.**
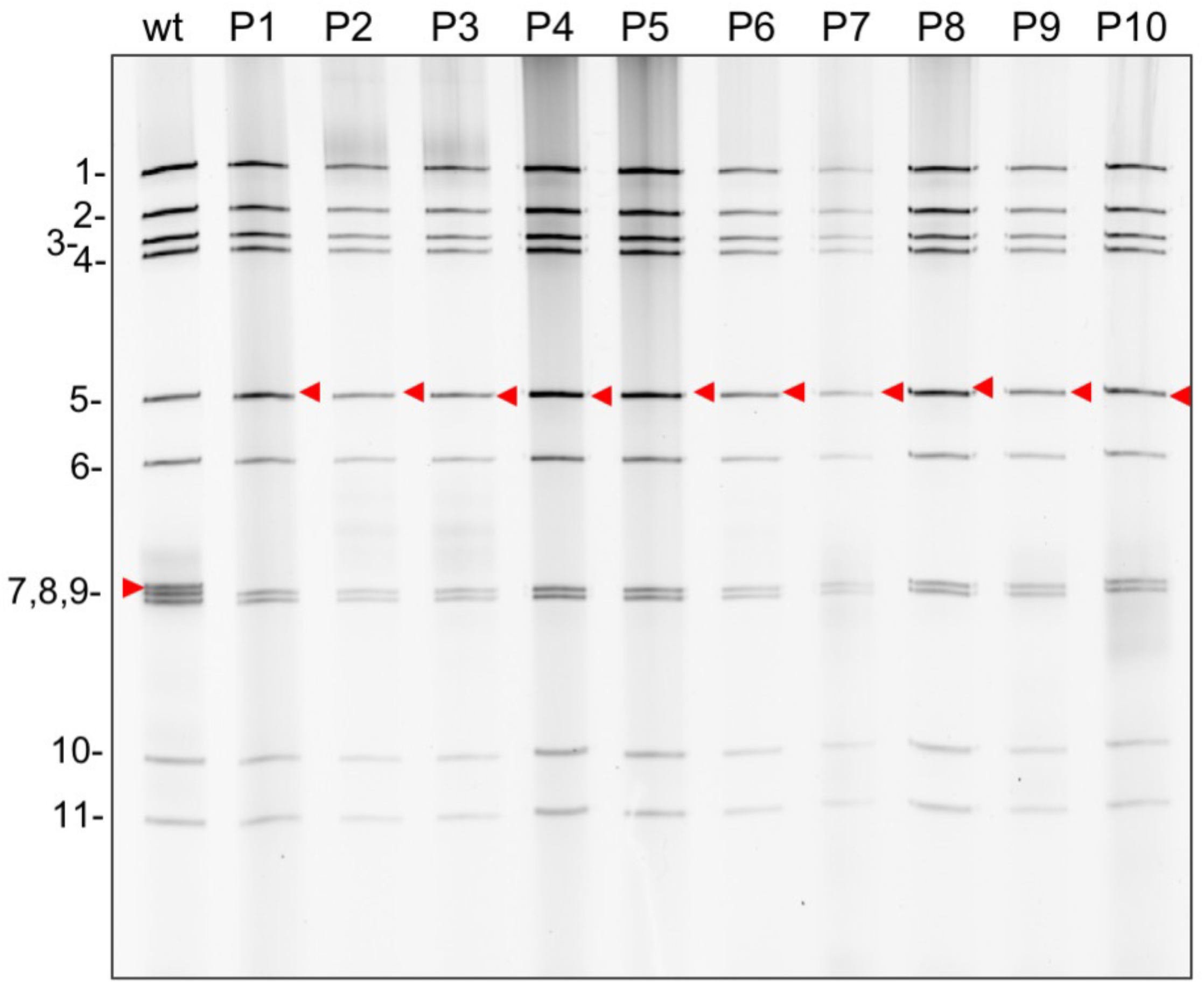
Genetic stability of rSA11/NSP3-FL-UnaG. Virus was serially passaged 10-times in MA104 cells using 1:1000 dilutions of infected cell lysate as inoculum. RNA was recovered from infected cell lysates using Trizol, treated 30 min with RNase T1 at room temperature, and resolved by electrophoresis on a 8% polyacrylamide gel. RNA gel was detected by staining gel with ethidium bromide. Red arrow denotes the position of the segment 7 (NSP3) RNA.

### Fluorescence signal of rSA11/NSP3-FL-UnaG

Examination of MA104 cells infected with rSA11/NSP3-FL-UnaG by live-cell imaging from 0 to 16 h p.i. confirmed that UnaG was functional, emitting fluorescent light in a range overlapping green fluorescent protein (GFP) (Fig. 6A, Movie S1) (Rodriguez et al., 2017). At early times of infection (0 to 8 h p.i.), the signal localized predominantly to the cytoplasm. At later times, UnaG fluorescent signal was also readily detected in the nucleus (Move S1). To contrast the intensity of the fluorescent signal produced by recombinant viruses expressing UnaG fused to NSP3 versus NSP1, we generated rSA11/NSP1-FL-UnaG (Fig. 7) using the optimized NP868R-based RG protocol. To produce this virus, a FL-UnaG ORF terminating with a stop codon was inserted into NSP1 ORF of the segment 5 cDNA of pT7/NSP1SA11. As illustrated in Fig. 7, pT7/NSP1-FL-UnaG produces a 2.1-kb RNA that encodes a 575-aa protein instead of the 1.6-kb RNA and 520-aa protein of pT7/NSP1SA11. The segment-5 protein product of pT7/NSP1-FL-UnaG ends with the same Flag-UnaG cassette as the segment-7 protein product of pT7/NSP3-FL-UnaG.

**Figure 6.**
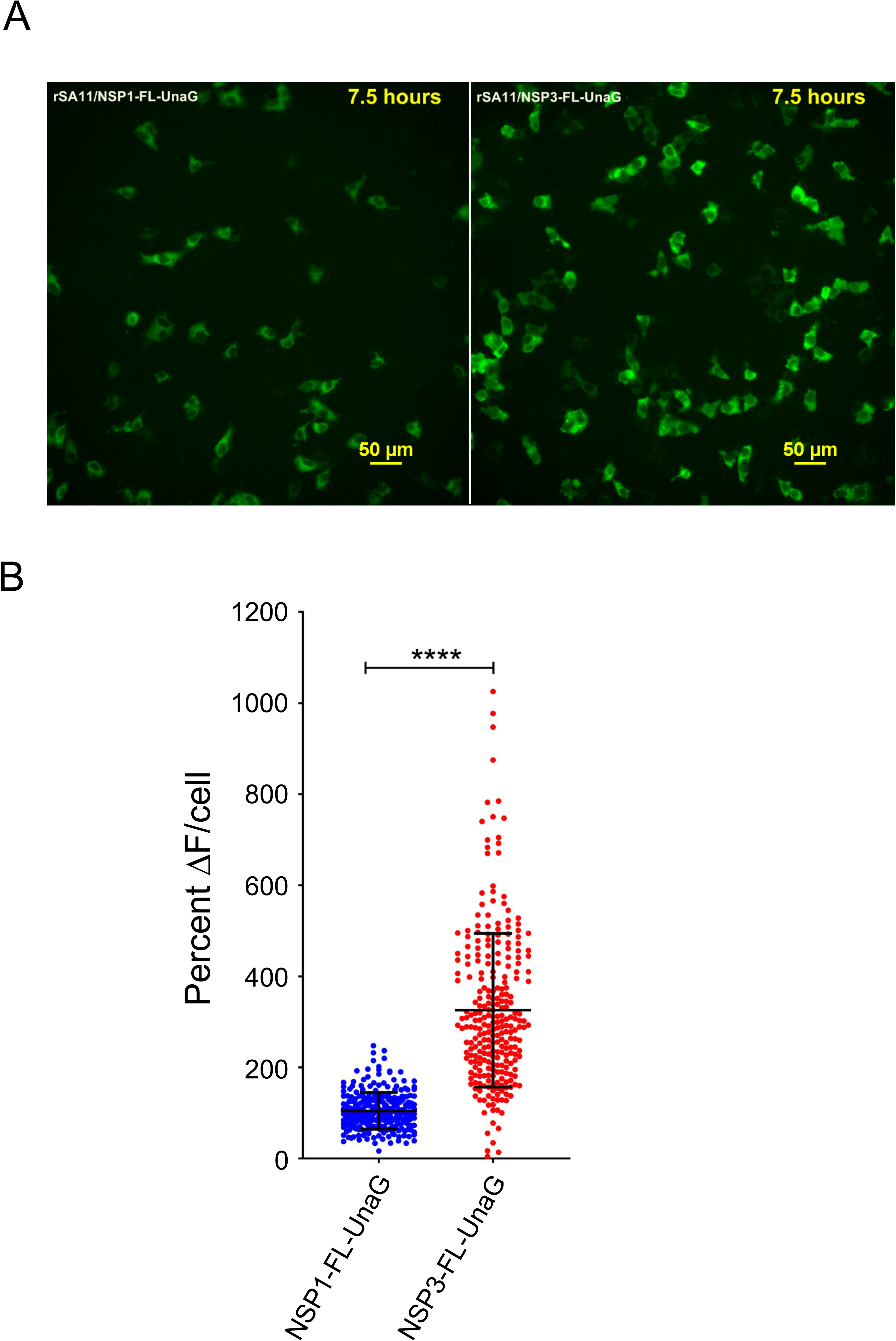
Comparison of UnaG expression by rSA11/NSP3-FL-UnaG versus rSA11/NSP1-FL-UnaG. (A) Images taken at 7.5 h p.i. with an epifluorescence Nikon TiE inverted microscope with a 20x Plan Apo (NA 0.75) differential interference contrast (DIC) objective. (B) Quantification of percent change in UnaG fluorescence (ΔF) in MA104 cells infected with rSA11/ NSP1-FL-UnaG or rSA11/NSP3-FL-UnaG. Single cell analysis (N=270 cells each) was performed to determine the percent ΔF from baseline. Data shown as mean ± standard deviation. ****p <0.0001

**Figure 7.**
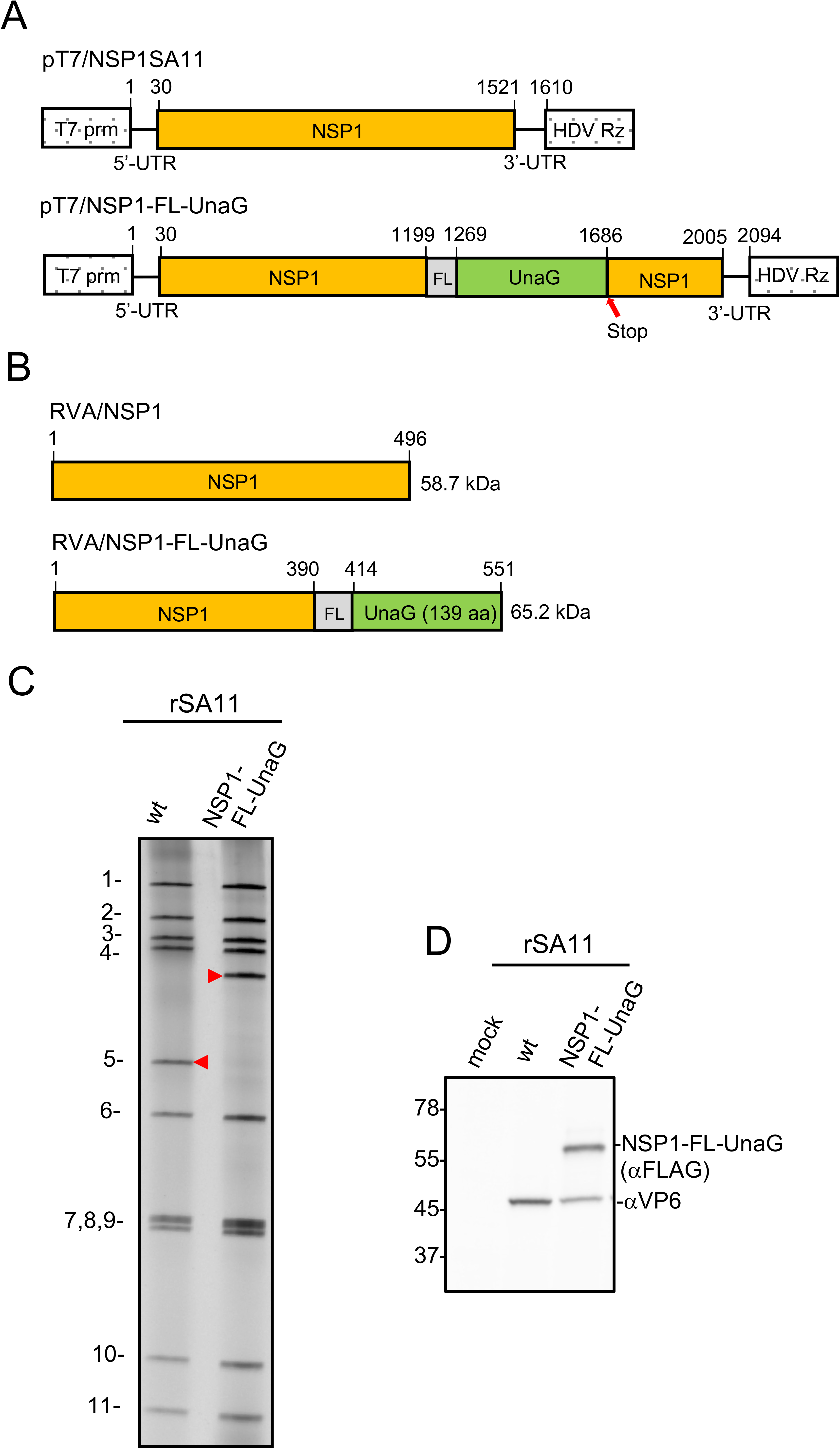
pT7 plasmids and rotaviruses expressing NSP1. (A) Organization of pT7 plasmids expressing wild type NSP1 and NSP1-FL-UnaG (+)RNAs, indicating locations of T7 promoter and Hepatitis delta virus self-cleaving ribozymes. The red arrow notes the position of the stop codon in the NSP1-FL-UnaG ORF. (B) Products of recombinant RVAs expressing wild type NSP1 and NSP1-FL-UnaG. (C) Profiles of the eleven genomic dsRNAs recovered from rSA11 viruses resolved by PAGE. Red arrow notes the position of the segment 5 dsRNA. (D) Western blot analysis of proteins present at 8 h p.i. in MA104 cells infected with recombinant viruses.

As determined by gel electrophoresis (Fig. 7) and sequencing (not shown), the genome of rSA11/NSP1-FL-UnaG included the expected large 2.1-kB segment 5 dsRNA. Immunoblot analysis of rSA11/NSP1-FL-UnaG-infected cell lysates using anti-Flag antibody indicated that the virus also encoded the NSP1-FL-UnaG product. Based on plaque assay, rSA11/NSP1-FL-UnaG grew to a peak titer (0.8 - 1.0 × 10_7_) that was ~3-fold less than rSA11/wt and produced plaques on MA104 cells that were ~2-fold smaller than rSA11/wt (Fig. 4) The small plaque phenotype of rSA11/NSP1-FL-NSP1 is similar to small plaque phenotype described before for RVAs encoding truncated or altered NSP1 proteins (Patton et al., 2001).

Quantification of UnaG fluorescence signals generated by MA104 cells infected with rSA11/NSP3-FL-UnaG were approximately 3-times greater than those generated by cells infected with rSA11/NSP1-FL-UnaG (Fig. 6B). This result suggests that RVAs expressing fused NSP3-FPs may be more sensitive probes of viral infection that RVAs expressing fused NSP1-FPs.

### Stem-loop structure in the segment 7 3’-UTR

rSA11/NSP3-FL-UnaG was generated by placing a nonviral 500-bp insert into the segment 7 dsRNA at the junction of the NSP3 ORF and 3’-UTR. Similar recombinant RVAs have been made by inserting nonviral sequences between the NSP2 ORF and 3’-UTR of the segment 8 dsRNA (Navarro et al., 2013). Thus, the junction between the viral ORF and 3’-UTR may represent a site well suited for the introduction of long foreign sequences into RVA genome segments. Interestingly, for RVAs with naturally occurring genome rearrangements, this is the same site in segment 5, 6, 7, 10 and 11 dsRNAs in which viral sequence duplications have been noted to initiate (Ballard et al., 1992; Shen et al., 1994; Gault et al., 2001; Patton et al., 2001; Arnold et al., 2012). The 3’-UTR contains multiple *cis*-acting signals important to rotavirus replication, including sequences that are recognized by the RVA RNA polymerase VP1 and the translation enhancer NSP3 (Deo et al., 2002; Tortorici et al., 2003). The fact that well-growing genetically-stable recombinant RVAs have been recovered in which a viral ORF has been separated from its 3’-UTR indicates that *cis*-acting signals in the 3’-UTR continue to function even though displaced linearly a long distance from the remaining viral sequence of the RNA.

In a previous study (Navarro et al., 2013), an *in silico* RNA folding analysis {http://rna.tbi.univie.ac.at/cgi-bin/RNAfold.cgi} was performed to probe how the insertion of sequence duplications and foreign sequences effected the predicted secondary structure of the mutant segment 8 (+)RNAs used in making recombinant RVAs. The results showed that despite extensive differences in the overall folding predictions of the mutant RNAs, in all cases their 5’- and 3’-UTRs interacted to form stable 5’-3’ panhandles. In addition, the predictions all revealed an identical stem-loop structure projecting from the 5’ side of the 5’-3 panhandle, formed by residues that are highly conserved among RVA segment 8 RNAs. The conservation of the structure and its sequence suggested that the stem-loop may function as a segment specific packaging signal (Navarro et al., 2013). We performed a similar *in silico* RNA folding analysis, contrasting the secondary structures predicted for the segment 7 RNAs of rSA11/wt and rSA11/NSP3-FL-UnaG. The results showed that the overall secondary structures predicted for the RNAs differed considerably, with the notable exception that extending from the 3’-UTR of both RNAs was a long (~70 base) stable stem-loop structure formed by sequences that are highly conserved in RVA segment 7 RNAs (Fig. 8). The stability and location of the stem-loop suggests that this structure may function as a segment specific packaging signal, in a manner previously proposed for the conserved stem-loop detected in the segment 8 RNA.

**Figure 8.**
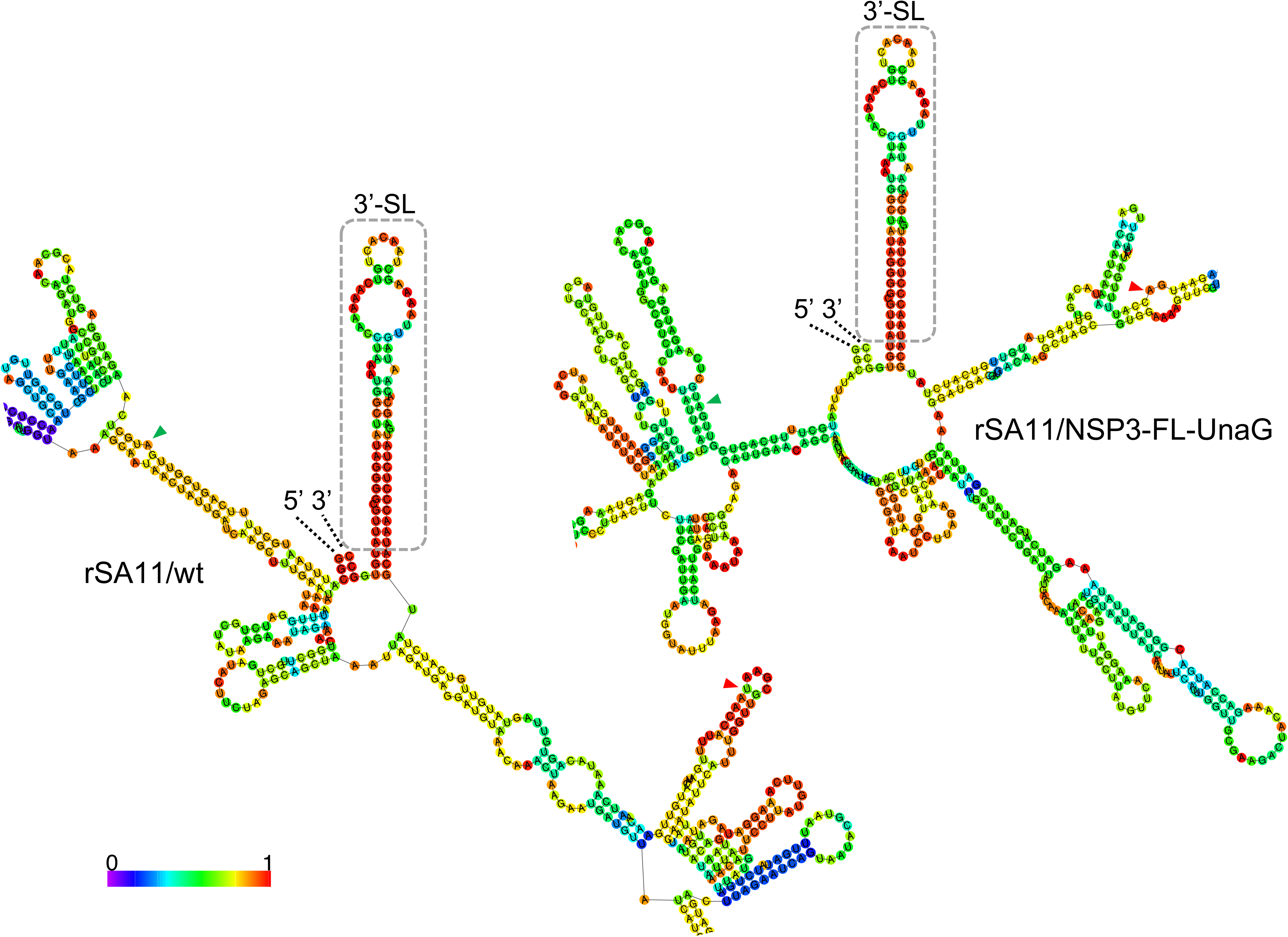
Conservation of a predicted stable stem-loop structure formed by the 3’-UTR sequence of rSA11/wt and rSA11/NSP3-FL-UnaG. Secondary structures associated with minimum free energy were calculated for segment 7 (+)RNAs using *RNAfold* {http://rna.tbi.univie.ac.at/} and color coded to indicate base-pairing probability (Hokacker, 2003; Hofacker et al., 1994). Portions of the secondary structures are shown that include the 5’ and 3’ ends of the (+)RNAs (labeled) and the conserved 3’ stem-loops (3’SL) (boxed). Also labeled are the start and stop codons (green and red arrowheads, respectively) of both the NSP3 and NSP3-FL-UnaG ORFs.

### Summary

rSA11/NSP3-FL-UnaG is the first recombinant RVA to be described with a modified segment 7 dsRNA. Segment 7 joins segments 4 (VP4) (Johne et al., 2015; Mohanty et al., 2017), 5 (NSP1) (Kanai et al., 2019), 8 (NSP2) (Trask et al., 2010; Navarro et al., 2013), and 11 (NSP5/NSP6) (Komoto et al., 2017) as targets altered by RG and represents only the second RVA segment to be used as a vector for FP expression. Our analysis of rSA11/NSP3-FL-UnaG indicates that it is possible to generate recombinant RVAs that express FPs through their fusion to the C-terminus of NSP3. Given that NSP3 is expressed at moderate levels in infected cells, RVAs expressing NSP3-based FPs may be more effective indicators of viral replication in live cell imagining experiments and other fluorescence-based assay systems than RVAs expressing NSP1-based FPs, since NSP1 is expressed at low levels *in vivo* (Martinez-Alvarez et al., 2013). Although several recombinant RVAs that express FPs have been described, rSA11/NSP3-FL-UnaG is unique among them in that none of its ORFs have been deleted or interrupted. Instead, the only impact on rSA11/NSP3-FL-UnaG was to fuse its NSP3 ORF to a FL-UnaG ORF. Importantly, although the NSP3 ORF in RVA strains is not naturally extended and does not encode NSP3 fused to a downstream protein, the NSP3 ORF of group C rotaviruses (RVCs) is extended, encoding an NSP3 protein that is fused to a 2A stop-start translational element (Donnelly et al., 2001) and double-stranded RNA binding protein (dsRBP) (James et al., 1999; Langland et al., 1994). Given that RVC segment 6 encodes an NSP3 fusion protein, it seems likely that the NSP3 fusion protein of rSA11/NSP3-FL-UnaG remains functional, even when fused to a downstream protein. Interestingly, despite repeated attempts, we were unsuccessful in generating recombinant RVAs using mutated pT7/NSP3SA11 plasmids in which the NSP3 ORF was interrupted through insertion of stop codons (data not shown). This result implies that NSP3 is essential for RVA replication or is required to generate recombinant viruses using the RG system.

Our results suggest that the RVA segment 7 RNA can be re-engineered to function as an expression vector of foreign proteins, without compromising the function of any of the viral ORFs. It remains unclear how much foreign sequence can be inserted into the segment 7 RNA, but our analysis so-far indicates that it is possible to generate well replicating viruses carrying >500 bp of extra sequence. Given the remarkable flexibility so far noted in the ability of the rotavirus to accommodate changes in the size and sequences of its RNA, the virus may turn out to be particularly empowering tool for unraveling the shared mechanisms used by the *Reoviridae* to package and replicate their genomes.

## Acknowledgments

We are grateful to all the members of the Patton and Danthi laboratory for their support and encouragement on this project. Our thanks also go to Ulla Buckholtz and Peter Collins, NIAID, NIH for their gift of BHK-T7 cells. This work was supported by National Institutes of Health grants R03 AI131072 and R21 AI144881, Indiana University Start-Up Funding, and the Lawrence M. Blatt Endowment.

---

**Movie S1.** Expression of fluorescent signal by rSA11/NSP3-FL-UnaG and rSA11/NSP1-FL-UnaG over the course of infection. Images were collected at 6 min intervals from 0 to 16 h p.i. using an epifluorescence Nikon TiE inverted microscope with a 20x Plan Apo (NA 0.75) differential interference contrast (DIC) objective.

